# An explainable language model for antibody specificity prediction using curated influenza hemagglutinin antibodies

**DOI:** 10.1101/2023.09.11.557288

**Authors:** Yiquan Wang, Huibin Lv, Ruipeng Lei, Yuen-Hei Yeung, Ivana R. Shen, Danbi Choi, Qi Wen Teo, Timothy J.C. Tan, Akshita B. Gopal, Xin Chen, Claire S. Graham, Nicholas C. Wu

## Abstract

Despite decades of antibody research, it remains challenging to predict the specificity of an antibody solely based on its sequence. Two major obstacles are the lack of appropriate models and inaccessibility of datasets for model training. In this study, we curated a dataset of >5,000 influenza hemagglutinin (HA) antibodies by mining research publications and patents, which revealed many distinct sequence features between antibodies to HA head and stem domains. We then leveraged this dataset to develop a lightweight memory B cell language model (mBLM) for sequence-based antibody specificity prediction. Model explainability analysis showed that mBLM captured key sequence motifs of HA stem antibodies. Additionally, by applying mBLM to HA antibodies with unknown epitopes, we discovered and experimentally validated many HA stem antibodies. Overall, this study not only advances our molecular understanding of antibody response to influenza virus, but also provides an invaluable resource for applying deep learning to antibody research.

## INTRODUCTION

Discovery and characterization of monoclonal antibodies are central to the understanding of human immune response, as well as design of vaccines and therapeutics [1, 2]. As exemplified by SARS-CoV-2 research in the past few years, antibody discovery has dramatically accelerated due to the technological advancements in single-cell high-throughput screen [3] and paired B cell receptor sequencing [4]. Nevertheless, epitope mapping remains a major bottleneck of antibody characterization, which often involves the determination of individual antigen-antibody complex structures using X-ray crystallography or cryogenic electron microscopy (cryo-EM). As a result, there is a huge interest in developing methods for antibody specificity prediction.

Despite the huge diversity of human antibody repertoire with at least 10^15^ antibody sequences [5, 6], antibody responses from different individuals often utilize recurring sequence features to target a given epitope [7–15]. This phenomenon is also known as convergent or public antibody response. Traditionally, antibody specificity prediction has mainly relied on biophysical models [16]. However, the observation of public antibody response suggests that antibody specificity prediction can also be achieved by an orthogonal, data driven approach. Specifically, with a sufficiently large sequence dataset of human antibodies that share a common epitope, a purely sequence-based model can be trained to predict whether an antibody targets this given epitope or not.

Recently, the application of natural language processing has revolutionized protein structure and function prediction as well as protein design [17–23]. While several language models for antibodies have also been developed [24–26], none of them enables antibody specificity prediction to the best of our knowledge. One of the major barriers to developing a language model for antibody specificity prediction is the lack of systematically assembled datasets for model training, which would require both sequence and epitope information for individual antibodies. Although many studies have reported sequences of antibodies with known epitopes, such information is often not centralized. Database such as CoV-AbDab, which documents the sequence and epitope information for >10,000 antibodies to coronavirus [27], is absent for most pathogens including influenza virus.

Hemagglutinin (HA) is the major antigen of influenza virus and has a hypervariable globular head domain atop a highly conserved stem domain [28]. In this study, we manually curated 5,561 human antibodies to influenza hemagglutinin (HA) protein from research publications and patents. Recurring sequence features among these HA antibodies were identified, many of which were previously unknown. Using this dataset, we further developed a memory B cell language model (mBLM) for antibody specificity prediction based on seven specificity categories, including HA head and stem domains. Saliency map explanation of mBLM revealed that key binding motifs were learned during specificity prediction. Moreover, we successfully applied mBLM to discover HA stem antibodies with subsequent experimental validation.

## RESULTS

### A large-scale collection of influenza antibody information

We compiled a list of 5,561 human monoclonal antibodies to influenza HA from 60 research publications and three patents (**Table S1**). Information on germline gene usage, sequence, binding specificity (e.g. group 1, group 2, type A or B, etc.), epitope (head or stem), and donor status (e.g., infected patient, vaccinee, etc.), if available, was collected for individual antibodies. Among these antibodies, which were isolated from 132 different donors, 564 (10.1%) bind to the globular head domain and 518 (9.3%) bind to the stem domain. Epitope information was not available for the remaining 4,479 HA antibodies.

### HA head and stem antibodies have distinct sequence features

We first aimed to analyze this large dataset to examine the recurring sequence features of human antibody responses to influenza HA. Our analysis captured previously known germline gene preference for HA stem antibodies, such as IGHV1-69 [8, 29] and IGHD3-9 [7], as well as for HA head antibodies, such as IGHV2-70 and IGHD4-17 (**Figure 1A**, **Figure 1C, and Figure S1**) [30]. Other recurring sequence features were also observed in our analysis, such as the enrichment of IGKV3-11, IGKV3-15, and IGKV3-20 among HA stem antibodies, as well as IGKV1-33 and IGLV3-9 among HA head antibodies (**Figure 1B**). In addition, our analysis discovered five public clonotypes that target influenza type B HA (clonotypes 13, 16, 56, 89, and 117) that have not been described previously to the best of our knowledge (**Figure S2 and Table S1**).

**Figure 1.**
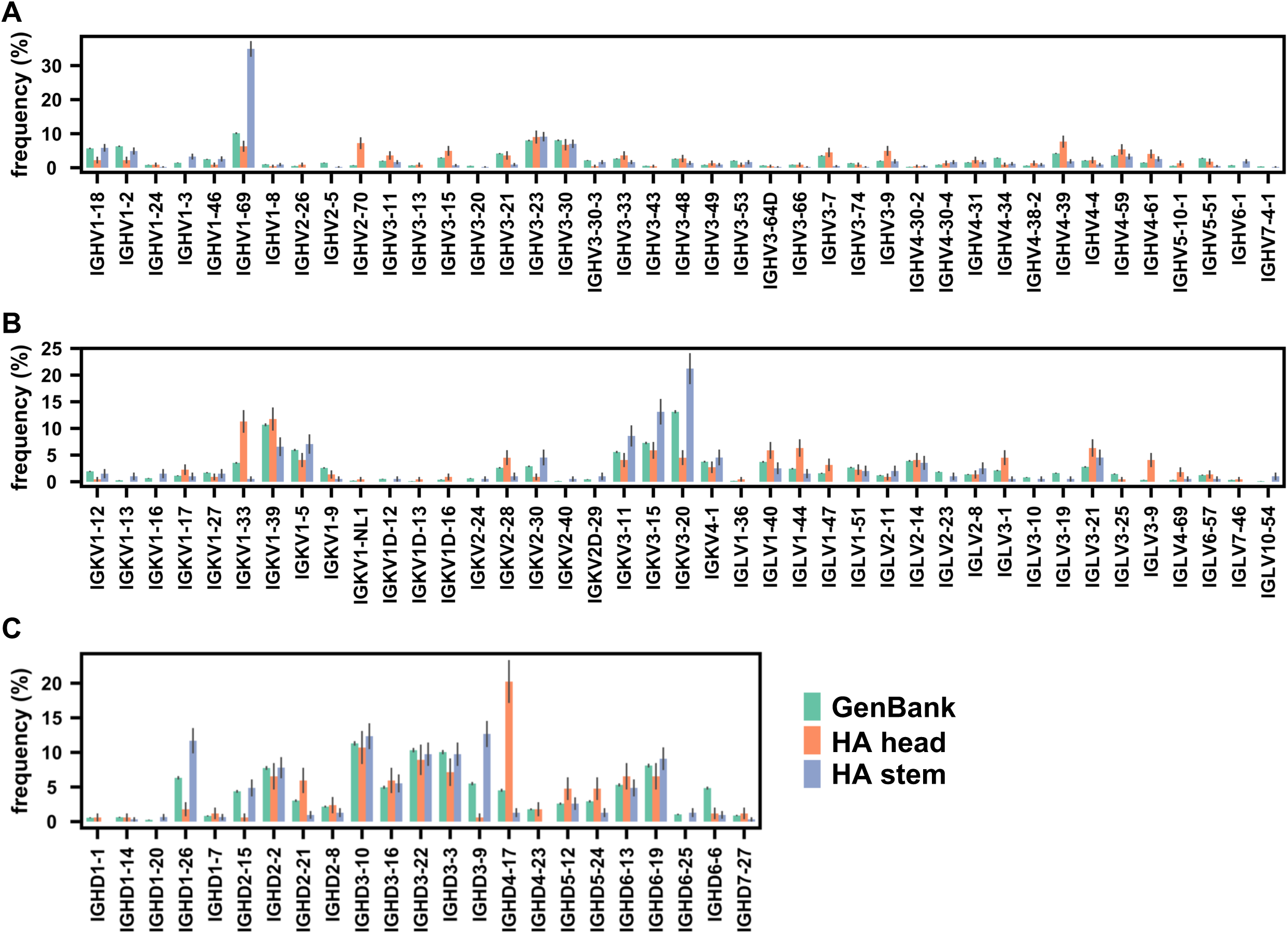
Germline gene usages in influenza HA antibodies. **(A)** The IGHV gene usage, **(B)** IGK(L)V gene usage, and **(C)** IGHD gene usage in antibodies to HA head domain (orange) and HA stem domain (blue). For comparison, germline gene usages of all antibodies from Genbank are also shown (green). To avoid being confounded by B-cell clonal expansion, a single clonotype from the same donor is considered as one antibody (**see Methods**).

The high prevalence of IGHD4-17 among HA head antibodies stood out to us. It is known that the second reading frame of IGHD4-17 encodes a YGD motif (**Figure S3A**) and can pair with IGHV2-70 to form a multidonor antibody class targeting the receptor binding site in the HA head domain [30]. However, our analysis here demonstrated that IGHD4-17 could pair with other IGHV genes to target diverse epitopes in the HA head domain (**Figure S3B, Figure S4A, and Table S2**). Most of these antibodies contain an IGHD4-17-encoded YGD motif in the complementarity determining region (CDR) H3 (**Table S2**). Consistently, CDR H3 with a YGD motif was observed in 12.8% of the HA head antibodies, but only in 0.8% and 2.0% of the HA stem antibodies and all antibodies from GenBank (**Figure S4B and Table S3**), respectively. These observations suggest the versatility of the IGHD4-17-encoded YGD motif in targeting multiple epitopes in the HA head domain, similar to the ability of IGHV3-53 to engage different epitopes in SARS-CoV-2 spike (S) receptor-binding domain (RBD) [31, 32].

While the major antigenic sites in the HA head domain largely consist of hydrophilic and charged amino acids [33–36], HA stem antibodies are known to commonly target a hydrophobic groove [37]. Consistently, the CDR H3 sequences of HA stem antibodies had significantly higher hydrophobicity than those of HA head antibodies (*p* = 0.001) (**Figure 2A**). Such difference was more pronounced when we only considered the tip of the CDR H3, which locates in the center of the CDR H3 sequence and is typically important for binding (*p* = 4e-12) (**Figure 2B**). In contrast, the CDR H3 lengths of antibodies to HA head and stem domains did not differ significantly (*p* = 0.38) (**Figure 2C**). Overall, these analyses reveal distinct recurring sequence features between HA head and stem antibodies.

**Figure 2.**
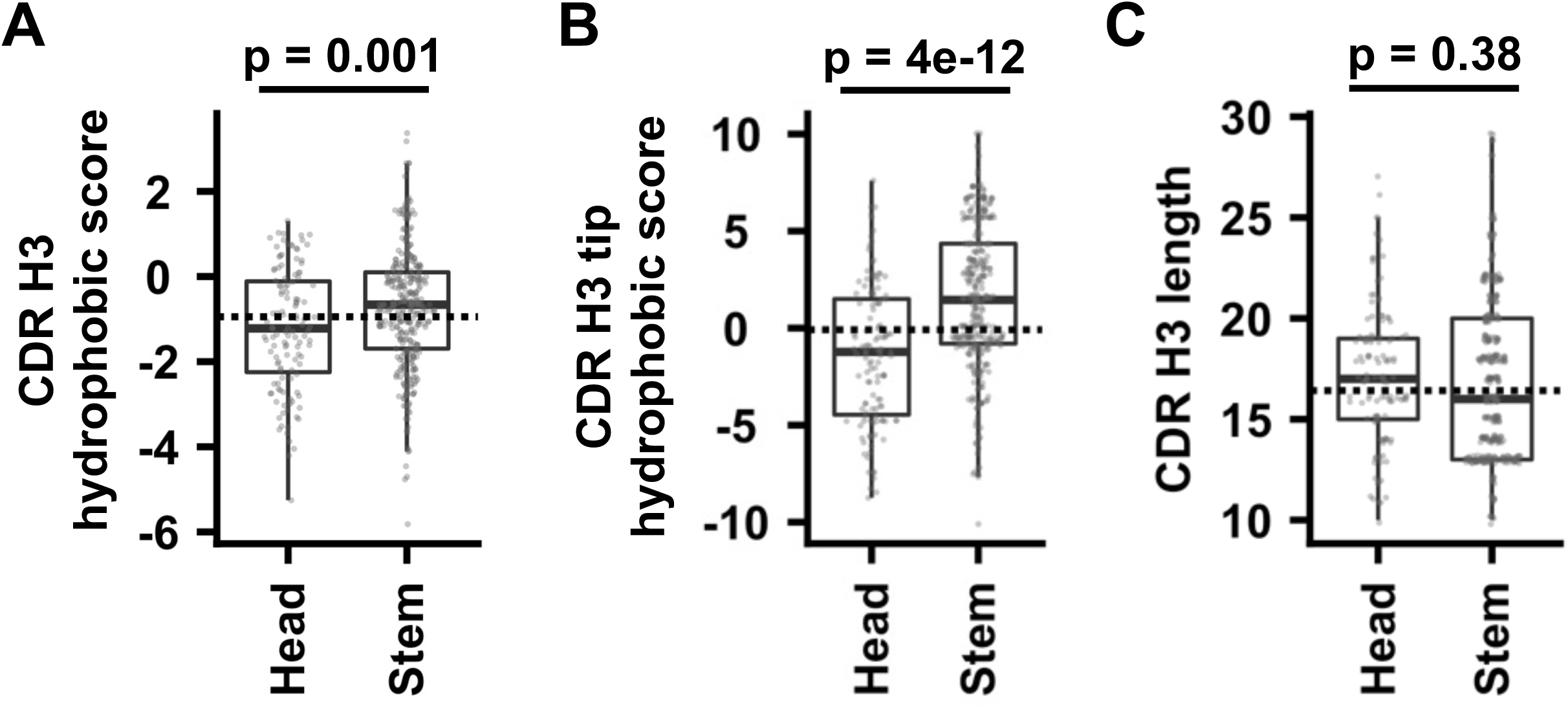
Hydrophobicity of CDR H3 sequences. **(A-B)** The hydrophobicity scores of **(A)** CDR H3 and **(B)** CDR H3 tip, as well as **(C)** the CDR H3 length are compared between antibodies to HA head and HA stem domains. The p-values were computed by two-tailed Student’s t-tests. For the boxplot, the middle horizontal line represents the median. The lower and upper hinges represent the first and third quartiles, respectively. The upper whisker extends to the highest data point within 1.5x inter-quartile range (IQR) of the third quartile, whereas the lower whisker extends to the lowest data point within 1.5x IQR of the first quartile. Each data point represents one antibody. The horizontal dotted line indicates the mean among antibodies from Genbank.

### Antibody specificity prediction using mBLM

Our previous work has shown that antibodies with different specificities can be distinguished using a sequence-based machine learning model that has a simple architecture with one transformer encoder for each CDR, followed by a multi-layered perceptron (“CDR encoders”) [15]. Here, we postulated that a language model could offer better performance, given the recent success of applying language models to predict protein structures and functions [17–23]. Specifically, we aimed to pre-train a memory B cell language model (mBLM) to learn the intrinsic “grammar” of functional antibodies, and to subsequently distinguished between HA head and stem antibodies, as well as antibodies to other antigens.

Briefly, mBLM was pre-trained to predict masked amino acid residues in the context of paired heavy and light chain antibody sequences, using a total of 253,808 unique paired antibody sequences from GenBank [38] and Observed Antibody Space [39] (**see Methods**). For antibody specificity prediction, mBLM was fine-tuned by using the final-layer embeddings of the pre-trained mBLM, followed by a multi-head self-attention block and a multi-layer perceptron (MLP) block (**Figure 3A**). Our prediction was based on seven specificity categories, namely influenza HA head, influenza HA stem, HIV, SARS-CoV-2 S NTD, SARS-CoV-2 S RBD, SARS-CoV-2 S S2, and others (none of the above). Since many antibodies in these specificity categories did not have light chain sequence available, only heavy chain sequences were used for specificity prediction (**see Methods**). Of note, the highest pairwise sequence identity between the test and training sets was 80%. In other words, the pairwise sequence identity between individual antibody sequences in the test set and the training set was at least 20% (i.e. 26 amino acids). As indicated by the confusion matrix analysis and F1 score, mBLM had a decent performance on the test set (**Figure 3B-C**). The F1 score on the test set was 0.75 for mBLM, but only 0.49 for CDR encoders (**Figure 3C**). The performance of mBLM, which had 41 million parameters, was also slightly better than the pre-trained general protein language model ESM2 with 650 million parameters (F1 score on the test set = 0.74) [18]. This result demonstrates that mBLM is an efficient model for antibody specificity prediction.

**Figure 3.**
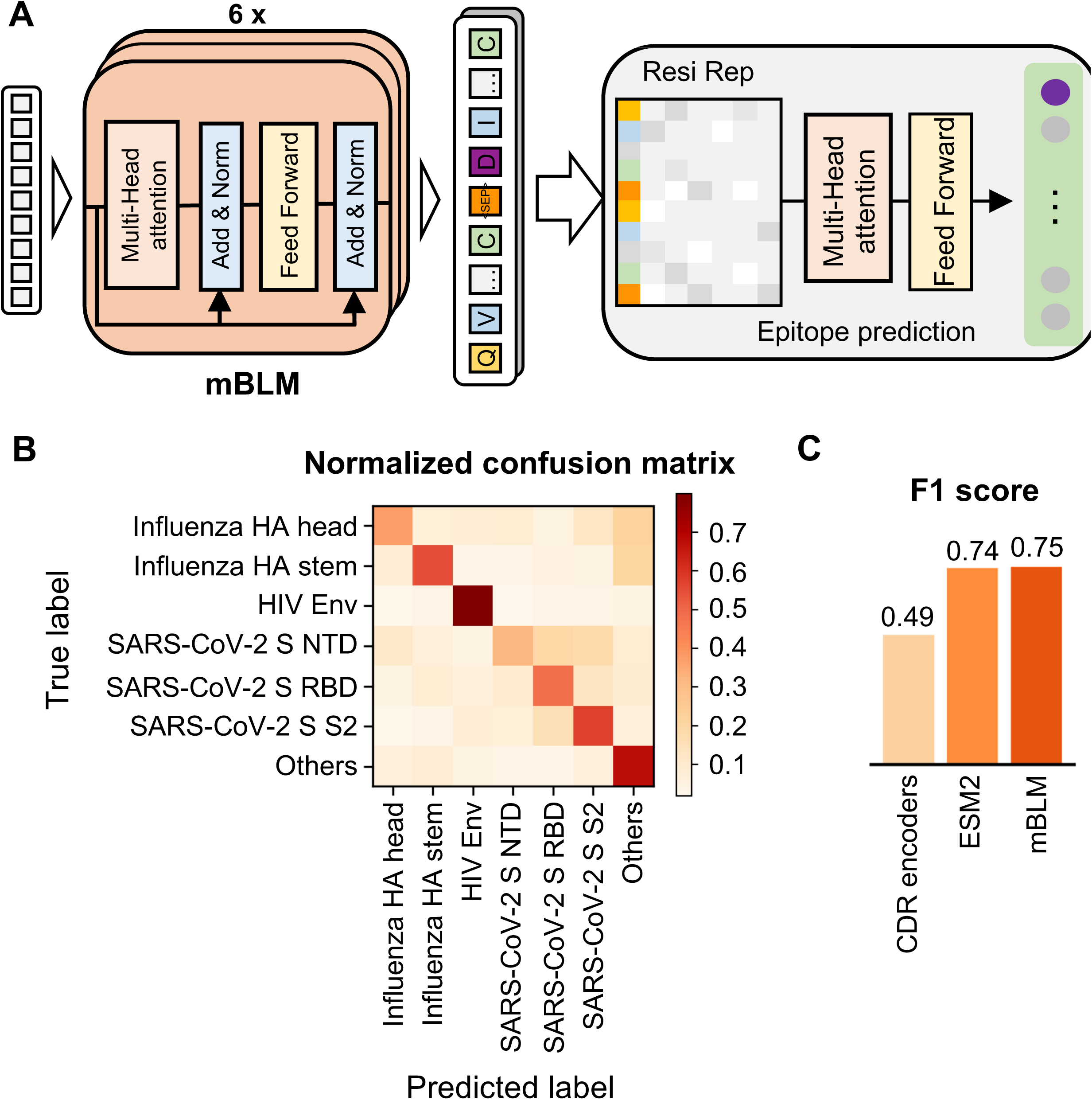
Antibody specificity prediction by memory B cell language model (mBLM). **(A)** Model architecture of mBLM is shown. Arrows indicate the information flow in the network from the language model to antibody specificity prediction, with a final output of specificity class probability. Resi Rep: residual level representation (i.e. the final-layer embeddings from pre-trained mBLM). **(B)** Model performance of mBLM on the test set was evaluated by a normalized confusion matrix. **(C)** The performance of different antibody specificity prediction models was evaluated by F1 score, which represents the weighted harmonic mean of the precision and recall. CDR encoders: our previous model using a transformer encoder to encode CDR sequences [15]. ESM2: a general protein language model [18].

### mBLM learned the sequence features of HA stem antibodies

Next, we aimed to understand what mBLM had learned for antibody specificity prediction. Recent advancements in the field of computer vision have employed Gradient-Weighted Class Activation Maps (Grad-CAMs) on CNN-based architectures to identify the determinants for classification decisions [40, 41]. Here, Grad-CAM was adopted to analyze the fine-tuned mBLM by quantifying the importance of individual amino acid residues for antibody specificity prediction. Our result indicates that residues with high importance, as indicated by the saliency score, were enriched in the CDRs (**Figure 4A**).

**Figure 4.**
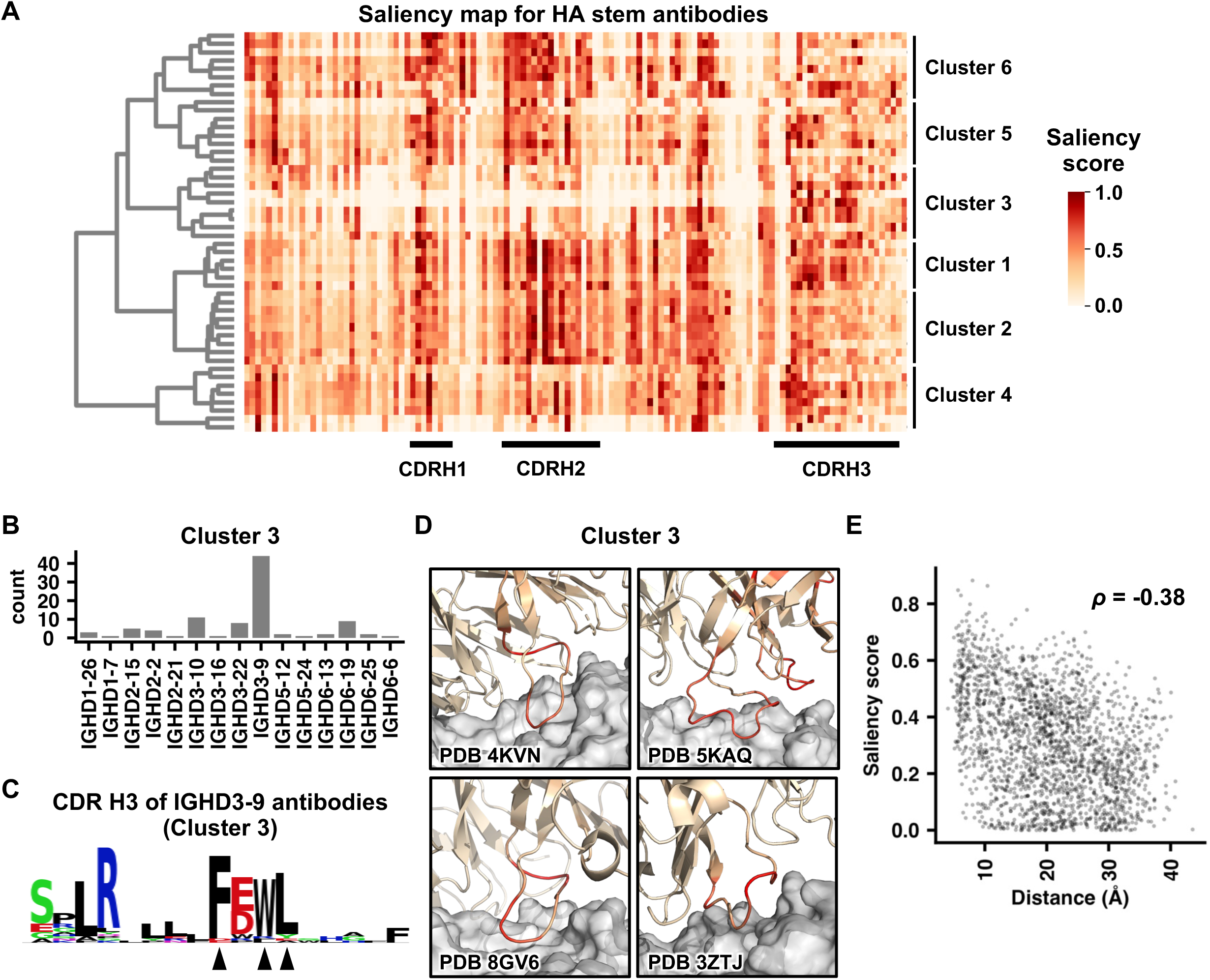
Explanation of mBLM using saliency score. **(A)** Saliency score for each residue in individual HA stem antibodies was shown as a heatmap. Each row represents a single HA stem antibody. X-axis represents the amino acid residue of the heavy chain. Regions corresponding to CDR H1, H2, and H3 are indicated. For visualization purpose, only 50 HA stem antibodies are shown. Six clusters of HA stem antibodies were identified using hierarchical clustering with Ward’s method. **(B)** IGHD gene usage among antibodies in cluster 3 is shown. **(C)** The saliency score of each CDR H3 residue in IGHD3-9 antibodies within cluster 3 was analyzed. The frequency of each amino acid for residues with a saliency score >0.5 is shown as a sequence logo. Arrows at the bottom indicate the residues of interest. **(D)** Saliency scores are projected on to the structures of four antibodies in cluster 3 (PDB 4KVN [49], PDB 5KAQ [42], PDB 8GV6 [54], and PDB 3ZTJ [47]). The color scheme is same as that in panel A. **(E)** The relationship between saliency score and distance to the antigen (i.e. HA stem) is shown as a scatter plot. Spearman’s rank correlation coefficient (*ρ*) is indicated. A total of 18 structures of HA stem antibodies in complex with HA were analyzed (PDB 3FKU, 3GBN, 3SDY, 3ZTJ, 4FQI, 4KVN, 4NM8, 4R8W, 5JW3, 5KAN, 5KAQ, 5K9K, 5K9O, 5K9Q, 5WKO, 6E3H, 6NZ7, and 8GV6) [7, 29, 42, 44–54].

Based on the saliency score pattern, we further identified six clusters of HA stem antibodies. These clusters captured several known sequence features of HA stem antibodies. For example, most antibodies in cluster 3 are encoded by IGHD3-9 (**Figure 4B**), which is known to be enriched among HA stem antibodies (**Figure 1C**) [7]. Among IGHD3-9 antibodies in cluster 3, we observed an FxWL motif in the CDR H3 with high saliency score (**Figure 4C**). As described previously, many IGHD3-9 antibodies are featured by a LxYFxWL motif in the CDR H3 [7]. Therefore, our result indicates that the fine-tuned mBLM partially learned a known CDR H3 motif for predicting HA stem antibodies. Other known sequence features of HA stem antibodies were also learned by mBLM, including IGHV1-18 with a QxxV motif in the CDR H3 (**Figure S5A-B**) [42], IGHV1-69 with Y98 (**Figure S5A-D**) [8], and IGHV6-1 with an FGV motif in the CDR H3 (**Figure S5E-F**) [43].

When we projected the saliency score of individual residues on the structures, residues closer to the epitope appeared to have a higher saliency score (**Figure 4D and Figure S5G-I**). Consistently, through systematically analyzing 18 structures of HA stem antibodies [7, 29, 42, 44–54], we found that the saliency score of individual residues in HA stem antibodies and their distance to HA exhibited a moderate negative correlation (Spearman’s rank correlation = -0.38, **Figure 4E**). Together, our result indicates that the fine-tuned mBLM could identify residues that were critical for binding and utilized them for specificity prediction, despite structural information was not used for model training.

To gain additional insights into the learned features of mBLM, we analyzed the final-layer embeddings of the pre-trained mBLM using t-SNE (t-distributed Stochastic Neighbor Embedding). Specifically, heavy chain sequences in the training set for fine-tuning were projected into a two-dimensional space according to the embeddings. The result showed clustering of antibodies that belonged to the same V gene family (**Figure S6A**). Moreover, antibodies from the same specificity category also tended to cluster together (**Figure S6B**). These observations demonstrated that even during the pre-training step, mBLM partially learned the sequence features that were determinants for antibody specificity, hence specificity prediction.

### Discovering HA stem antibodies using mBLM

There are two non-overlapping epitopes in the HA stem, namely central stem epitope [44, 45] and anchor stem epitope [55, 56]. A recent study has reported the isolation of 60 HA antibodies to the central stem epitope, and 38 to the anchor stem epitope [57]. While these antibodies were not in the HA antibody dataset that we assembled (**Table S1**), they provided an additional opportunity to test the fine-tuned mBLM. Among the 60 antibodies to the central stem epitope, the fine-tuned mBLM correctly predicted 67% (40/60) as HA stem antibodies (**Figure 5A**). In contrast, among the 38 antibodies to the anchor stem epitope, only 8% (3/38) were predicted as HA stem antibodies (**Figure 5A**). The poor performance of the fine-tuned mBLM on antibodies to anchor stem epitope was likely due to lack of antibodies to anchor stem epitope in the dataset that we assembled (**Table S1**). In fact, antibodies to anchor stem epitope have only been extensively characterized two years ago [56]. These results suggest that HA stem antibodies correctly predicted by mBLM would mostly target the central stem epitope.

**Figure 5.**
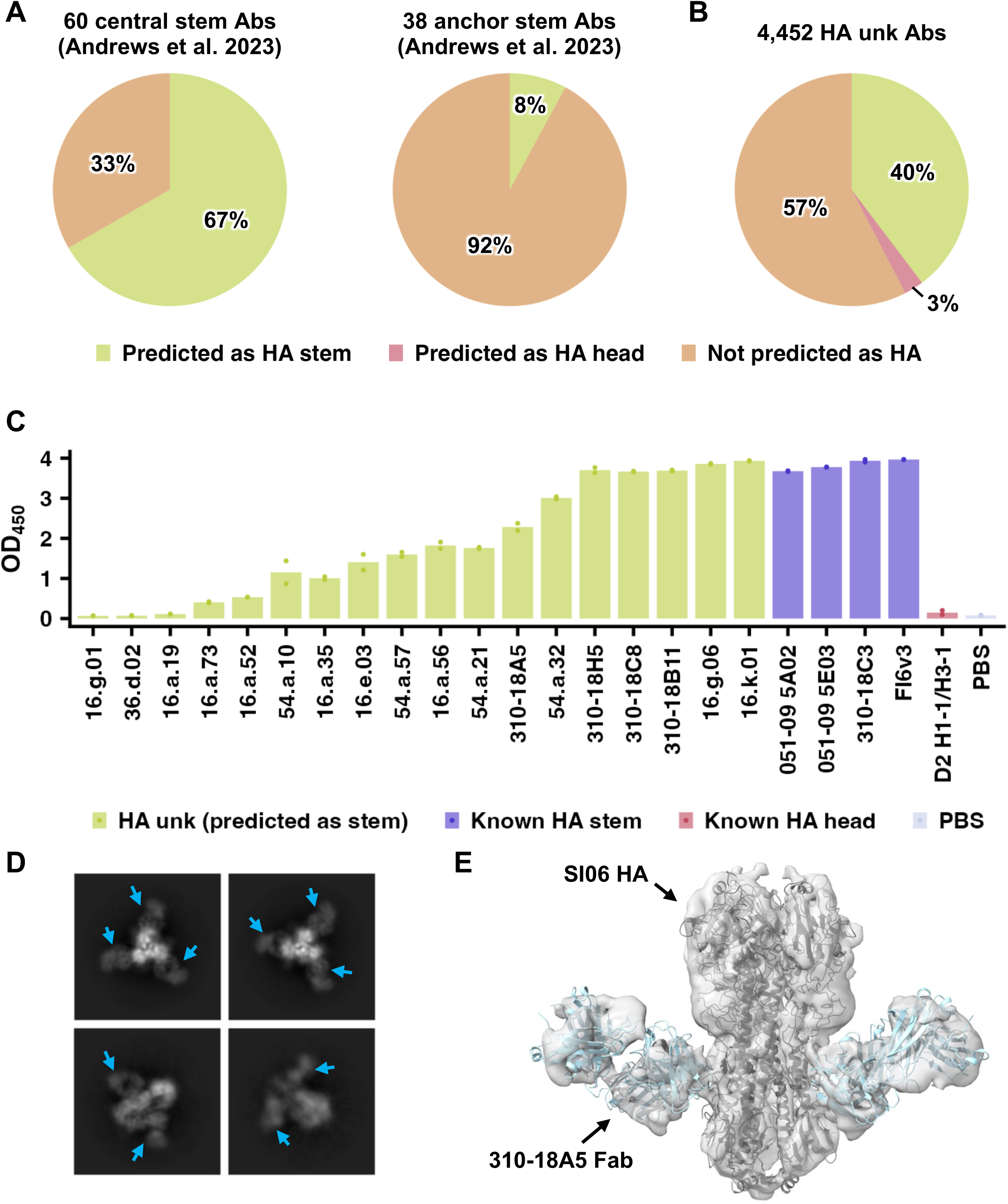
Discovery of HA stem antibody by mBLM. **(A-B)** mBLM was applied to predict the specificity of **(A)** 60 antibodies to central stem epitope (left panel) and 38 to anchor stem epitope (right panel) that were reported recently [57], as well as **(B)** 4,452 HA antibodies with unknown epitopes (HA unk) in the dataset that we assembled. The fraction of antibodies that were predicted to bind to HA stem domain (Predicted as HA stem), HA head domain (Predicted as HA head), or to other antigens (Not predicted as HA) is shown. **(C)** Using ELISA, the binding of 18 HA unk antibodies that were predicted as HA stem antibodies was tested against mini-HA, which is an H1 stem-based construct [58]. Four known HA stem antibodies (051-09 5A02, 051-09 5E03, 310-18C3, and FI6v3) [47, 63, 64] were included as positive control. D2 H1-1/H3-1, which is a known HA head antibody [65], was included as negative control. In this binding experiment, antibodies were not purified from the supernatant and thus their concentrations were unknown. **(D)** Representative 2D classes from cryo-EM analysis of 310-18A5 Fab in complex with H1N1 A/Solomon Islands/3/2006 (SI06) HA are shown. Cyan arrows point to the 310-18A5 Fabs. **(E)** Cryo-EM 3D reconstruction of 310-18A5 Fab in complex with SI06 HA. Structural models of SI06 HA (PDB 6XSK) [66] and CR9114 (PDB 4FQH) [48] were docked into the 3D reconstruction.

Among the 5,561 HA antibodies in the dataset that we assembled (**Table S1**), 80% (4,479/5,561) have unknown epitopes, of which 4,452 have heavy chain sequence information available. Subsequently, we applied the fine-tuned mBLM to predict the specificities of these 4,452 antibodies. While 40% (1,769/4,452) were predicted as HA stem antibodies, only 3% (119/4,452) were predicted as HA head antibodies (**Figure 5B**). HA head antibodies were expected to have a much higher sequence diversity than HA stem antibodies, because the HA head domain has a huge sequence diversity across influenza strains and subtypes, unlike the highly conserved HA stem domain [28]. Consequently, the poor performance of the fine-tuned mBLM on HA head antibodies was likely due to insufficient sequences of HA head antibodies in our training set.

To experimentally validate our prediction result, 18 antibodies that were predicted to target HA stem were individually expressed and tested for binding to mini-HA, which is an HA stem-based construct without the HA head domain [58]. Our enzyme-linked immunosorbent assay (ELISA) result showed that 83% (15/18) could bind to mini-HA (**Figure 5C**). The remaining 3 antibodies also exhibited binding to mini-HA when tested at a high concentration (**Figure S7A**). We further selected one of the validated HA stem antibodies, 310-18A5, for additional characterization. Biolayer interferometry indicated that 310-18A5 had a strong binding affinity against the HA from H1N1 A/Solomon Island/3/2006 (K_D_ = 0.2 nM, **Figure S7B**) as well as mini-HA (K_D_ = 1.0 nM, **Figure S7C**). Besides, 310-18A5 had neutralization activity against two antigenically distinct H1N1 strains (**Figure S7D**). Consistently, cryo-EM analysis confirmed that 310-18A5 bound to the HA stem domain (**Figure 5D-E and Table S4**). Overall, these results demonstrate that the fine-tuned mBLM enables discovery of antibodies to known epitopes.

## DISCUSSION

While influenza HA antibodies have been studied over decades, there has been a lack of effort to summarize the information about these antibodies. In this study, we performed a large-scale analysis of more than 5,000 influenza HA antibodies by mining research publications and patents. Although many recurring sequence features of influenza HA antibodies have previously been reported in individual studies [7, 8, 29, 30, 42, 43, 56], our results revealed additional ones that have not been described to the best of our knowledge. For example, our study discovered the enrichment of YGD motif in the CDR H3 of HA head antibodies as well as multiple public clonotypes to influenza type B HA. We further developed a language model for antibody specificity prediction, which was subsequently applied to discover HA stem antibodies. Overall, this work not only advances the molecular understanding of influenza HA antibodies, but also provides an important resource for the antibody research community (**Table S1**).

Discovering antibodies to a specific antigen of interest typically requires less efforts than epitope mapping. Consistently, epitope information (head or stem) is available for only ∼20% of HA antibodies in our dataset. Nevertheless, we were able to utilize these ∼20% of HA antibodies to train mBLM to identify HA stem antibodies among the remaining ∼80% with no epitope information. This result demonstrates that mBLM can accelerate epitope mapping. Although our work here applied mBLM to predict antibody specificity based on seven specificity categories, it can be fine-tuned to extend to any specificities as long as sufficient and diverse antibody sequences with such specificities are available. Given that many antibodies with different specificities have been characterized in the literature, future generalization of mBLM to additional antibody specificities will likely be achievable by extensive data mining (see discussion below). Besides, the continuous improvement of the speed of antibody discovery and characterization will also be beneficial, if not essential [3, 4].

The success of applying deep learning model to protein research can largely be attributed to the presence of databases such as Protein Data Bank (PDB) [59], UniProt [60], UniRef [61], which describe the sequence-structure-function relationships. Similarly, most, if not all, existing models for antibody specificity prediction were trained using structural information of antibody-antigen interactions in PDB [16]. Nevertheless, the epitopes of most antibodies in the literature are mapped by non-structural approaches, such as competition or mutagenesis experiments [62]. These epitope mapping data, despite being obtained by non-structural approaches, are tremendously useful for training a model for antibody specificity prediction as shown by our study here. Consequently, future efforts should focus on establishing a centralized database that describes the sequence-specificity relationship for antibodies, even for those without structural information available. Such database will allow the power of deep learning models to be fully harnessed in antibody research.

## ACKNOWLEDGEMENTS

This work was supported by National Institutes of Health (NIH) DP2 AT011966 (N.C.W.), R01 AI167910 (N.C.W.), the Michelson Prizes for Human Immunology and Vaccine Research (N.C.W.), and the Searle Scholars Program (N.C.W.). We thank Kristen Flatt at the UIUC Materials Research Laboratory Central Research Facilities for assistance with cryo-EM experiments, as well as Meng Yuan and Zongjun Mou for help discussion.

## AUTHOR CONTRIBUTIONS

All authors conceived and designed the study. Y.W., H.L., and N.C.W. assembled the dataset. Y.W., Y.H.Y., and N.C.W. performed data analysis. H.L., I.R.S., D.C., Q.W.T. T.J.C.T., A.B.G. performed the antibody binding experiments. R.L., C.S.G., and X.C. purified the proteins and performed the cryo-EM analysis. Y.W., H.L., R.L., and N.C.W. wrote the paper and all authors reviewed and/or edited the paper.

## DECLARATION OF INTERESTS

N.C.W. consults for HeliXon. The authors declare no other competing interests.

## METHODS

### Collections of antibody information

Sequences of each human monoclonal antibody were from the original papers and/or NCBI GenBank database (**Table S1 and Table S3**) [38]. For influenza HA antibodies, additional information, including binding specificity, donor IDs and PDB codes, was collected from the original papers (**Table S1**). Putative germline genes were identified by IgBLAST [67, 68]. Some studies isolated antibodies from multiple donors, but the donor identity for each antibody was not always clear. For example, some studies mixed B cells from multiple donors before isolating individual B cell clones. Since the donor identity could not be distinguished among those antibodies, we considered them from the same donor with “donors”, “vaccinees”, “patients”, or “cohorts” as the suffix of the donor ID. In addition, although two studies by Andrews et al. [69, 70] had shared donors from the same clinical trial (VRC 315, ClincialTrials.gov identifier NCT02206464), their antibody naming schemes were different. The IDs for these donors had a prefix “315” as described in the first study [69]. While the prefixes of antibody names from the first study matched the donor ID (e.g. antibody 315-02-1F07 was from donor 315-02) [69], some antibody names from the second study did not (e.g. antibody name with prefix “20A-518-30”) [70]. As a result, we assigned the donor ID to the antibodies from the second study by CDR H3 clustering. For example, since all CDR H3 clusters that contained antibodies with prefix 20A-605-30 also contained antibodies from 315-02, antibodies with prefix 20A-605-30 were assigned with a donor ID of 315-02.

### Identification of public clonotype

Using a deterministic clustering approach, CDR H3 sequences that had the same length and at least 80% amino acid sequence identity were assigned to the same CDR H3 cluster. As a result, CDR H3 of every antibody in a CDR H3 cluster would have >20% difference in amino acid sequence identity with that of every antibody in another CDR H3 cluster. A clonotype was defined as antibodies that shared the same IGHV/IGK(L)V genes with CDR H3s from the same CDR H3 cluster. A public clonotype was defined as a clonotype with antibodies from at least two donors. The epitope of each public clonotype was defined by its members.

### Germline gene usage analysis

To avoid being confounded by B-cell clonal expansion, a single clonotype from the same donor was considered as one antibody that represented the consensus sequence of the given clonotype. While all antibodies within a clonotype had the same IGHV/IGK(L)V genes (see above), they may not have the same IGHD gene, often due to ambiguity in IGHD-gene assignment by IgBlast. For germline gene usage analysis, the most common IGHD gene within a clonotype from the same donor was considered.

### Hydrophobic score of CDR H3

The hydrophobic score for a CDR H3 with a length n was computed as follow:

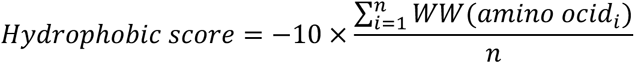

where WW represents the Wimley-White whole residue hydrophobicity scale [71] and amino acid_i_ represents the amino acid at position i. A higher hydrophobic score represents higher hydrophobicity. If the CDR H3 had an odd number of residues, the CDR H3 tip was defined as the three residues at the center of the CDR H3 sequence. If the CDR H3 had an even number of residues, the CDR H3 tip was defined as the four residues at the center of the CDR H3 sequence. The hydrophobic score of CDR H3 tip was computed in the same manner as that of CDR H3. To avoid being confounded by B-cell clonal expansion, a single clonotype from the same donor is considered as one antibody, in which the CDR H3 sequence represented the consensus among all members in the given clonotype.

### Datasets for model pre-training

A total of 267,871 paired antibody sequences from memory B cell sequencing data were downloaded from Observed Antibody Space database (BType = Memory-B-Cells) [39]. In addition, 12,487 paired antibody sequences were downloaded from NCBI GenBank database [38]. These antibody sequences were compiled into a single dataset and deduplicated by 95% sequence identity threshold. The deduplicated dataset was then partitioned into training (n = 229,773), validation (n = 15,375) and test sets (n = 8,660). The test set was generated by random sampling with different levels of maximum sequence identity to the training set (50%, 60%, 70%, 80%, and 90%), allowing robust evaluation of model performance. Of note, 90% maximum sequence identity indicated that none of the antibody sequences in the test set had >90% sequence identity with any of the sequences in the training set. In other words, the highest pairwise sequence identity between the test and training sets was 90%. To generate a balanced and robust training set, we implemented an upsampling technique based on the IGK(L)V genes. Specifically, we identified IGK(L)V genes with less than 5,000 counts and then performed random sampling to augment the dataset, ensuring each of these IGK(L)V genes had precisely 5,000 sequences. After upsampling, our training set had 467,018 paired antibody sequences. Of note, upsampling only applied to the training set, but not the validation and test sets.

### Sequences of antibodies with known specificities for model fine-tuning

Sequences of antibodies to “HA:Head” (influenza HA head) and “HA:Stem” (influenza HA stem) were from the curated dataset in this present study. Sequences of antibodies to “S:NTD” (SARS-CoV-2 spike NTD), “S:RBD” (SARS-CoV-2 spike RBD), and “S:S2” (SARS-CoV-2 spike S2) were from our previous study [15]. Sequences of antibodies to “HIV” (human immunodeficiency virus) and “Others” (none of the above) were collected from NCBI GenBank database [38]. Antibodies to “HIV” were classified as those from GenBank with the word “HIV” in the “References” or “Description” fields. Here, only heavy chain variable domain sequences were used for model fine-tuning. We performed sequence clustering with varying sequence identity cutoff (50%, 60%, 70%, 80%, 90%, and 95%) using cd-hit (-M 32000 -d 0 -T 8 -n 5 -aL 0.8 -s 0.95 -uS 0.2 -sc 1 -sf 1) [72]. We observed that at a cutoff of 90% sequence identity, sequences of antibodies with different specificities could be found within the same cluster, indicating that a stringent sequence identity cutoff of >90% was needed for accurate specificity prediction by traditional sequence clustering method. Based on this result, antibodies with unknown specificities, but shared >90% sequence identity with any antibody that belonged to “HA:Head”, “HA:Stem”, “HIV”, “S:NTD”, “S:RBD”, or “S:S2”, were discarded and not assigned to the “Others” category. Our final dataset for model fine-tuning contained the heavy chain sequences from a total of 388 antibodies to “HA:Head”, 509 antibodies to “HA:Stem”, 6,995 antibodies to “HIV”, 399 antibodies to “S:NTD”, 4112 antibodies to “S:RBD”, 682 antibodies to “S:S2”, and 15,043 antibodies to “Others”. This dataset was then partitioned into training, validation and test sets, with an approximate ratio of 8:1:1. To test model generalization, the test set was generated with a maximum sequence identity of 80% to the training set. In other words, the pairwise sequence identity between individual antibody sequences in the test set and the training set was at least 20% (i.e. 26 amino acids). We also applied the upsampling technique to the training set to ensure the number of antibody sequences in different specificity categories was balanced.

### Pre-trained memory B cell language model (mBLM)

#### Masked Language Modeling (MLM)

Masked language modeling such as Bidirectional Encoder Representations from Transformers (BERT) [73] has been shown as a powerful pretraining technique for language models, enabling contextual information to be captured and generalized to various downstream tasks. Here, mBLM was trained to predict the masked amino acids of input sequence based on surrounding context:

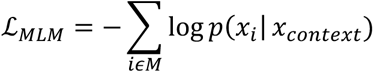

where *M* represents a randomly generated mask that includes 15% of positions *i* in the sequence

*x_i_*. The model was tasked with predicting the identity of the amino acids *x_i_* in the mask from the surrounding context *x*_()“*+,*_. Being trained to predict masked tokens, mBLM learned to understand the relationships between amino acid residues in a sequence, leading to a robust and effective language representations.

#### mBLM architecture

We adapted RoBERTa [74] as the basic model architecture, with the following hyperparameters: Tokenizer: ESM2 [18]

Token length: 150
Number of Layers: 6
Number of Attention heads: 12 Embedding dimension: 768
Feed-Forward Hidden Size: 3072
Dropout: 0.1

#### mBLM pre-training

mBLM was pre-trained with a context size of 250 tokens, which represented the amino acid sequences of both heavy and light chain variable domains. Since the total length of heavy chain and light chain variable domains was generally less than 250 amino acids, separation tokens were added in between. We adapted tokenizer from ESM2 [18], which converted amino acids into numerical representations (a total of 33 tokens including special tokens like [MASK]). The model was trained by masked language modeling (MLM) as described above. The model was optimized using Adam with *β*_1_ = 0.9, *β*_2_ = 0.999, *∈* = 10−8, and a learning rate of 5e-05. The model was trained using Huggingface transformers toolkit and efficiently distributed across one NVIDIA A100 and three NVIDIA RTX A5000. The entire pre-training process was completed within 24 hours, showcasing the efficiency and scalability of our approach.

### Model fine-tuning for specificity prediction

#### Model details

The final-layer embeddings from the pre-trained mBLM were extracted as the initial hidden state for the specificity prediction model. This initial state was then fed through a multi-head self-attention block and a multi-layer perceptron (MLP) block. An attention block was incorporated between the mBLM embeddings and the MLP significantly to enhance model interpretability. Within the attention block, the self-attention layer was followed by a layer normalization to normalize the output. Subsequently, an adaptive average pooling was applied to the attended representation to aggregate information across sequence dimension, resulting in a fixed size tensor with a shape that was defined by batch size and hidden dimension. The flattened tensor was then passed through the MLP block, comprising a series of fully connected layers, ReLU activation functions, and dropout operations. These layers transformed the high-dimensional representation to low-dimensional features. Finally, the output was passed through a fully connected layer with seven output units, each represented one of the seven specificity categories.

#### 15-fold cross-validation

To evaluate the robustness of our mBLM for specificity prediction, we employed a 15-fold cross-validation approach for fine-tuning, specificity inference, and model explanation. We randomly down/upsampled and split the data 15 times, resulting in a diverse set of sequences in each iteration. Then, the model underwent 15 rounds of training and testing. For each iteration, model performance was evaluated. The overall model performance was quantified as the average across all iterations. The final predicted class represented the mode.

#### mBLM fine-tuning

The model was trained using the PyTorch Lightning framework using Adam optimizer with a learning rate of 2e-05 and a batch size of 32. Early stopping was applied to monitor the validation loss.

#### ESM2 fine-tuning

Similar to mBLM fine-tuning, the final representations of ESM2 model (33 layers and 650 million parameters) were extracted as the initial hidden state for specificity prediction. This initial state was then fed through the attention and MLP blocks. The model was trained using the PyTorch Lightning framework using Adam optimizer with a learning rate of 1e-04 and a batch size 32. Early stopping was applied to monitor the validation loss. The best model checkpoint was saved.

#### Performance Metrics

The fine-tuned model was evaluated using the average F1 score, which represents the weighted harmonic mean of the precision and recall, as well as confusion matrix. The calculations were conducted using sklearn metrics functions [75].

### Model Interpretation

#### Gradient-weighted Class Activation Mapping (Grad-CAM) analysis

Grad-CAM, which is a class-discriminative localization technique that provides visual explanations for predictions made by CNN-based models [40], was used to identify residues in a protein sequence that are important for the prediction of a particular function [41]. To calculate Grad-CAM, we first computed the importance weights 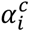 for the input sequence:

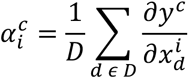

where 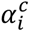 represents the global average pooling over embedding dimension *D* for the importance weights of residue *i* for predicting specificity class *c*. Then, the saliency map was obtained in a residue space by generating the weighted forward activation maps Α*^i^* , followed by a *ReLU* function:

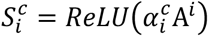

where 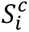 represents the relative importance (saliency score) of residue *i* to specificity class *c*. The *ReLU* function ensured that only features with positive influence on the functional label were preserved.

#### Saliency map clustering

We applied hierarchical clustering with Ward’s method to perform saliency map clustering. Euclidean distance was used to calculate the distance matrix that quantified the pairwise dissimilarity between saliency maps. We then used the linkage function to define the hierarchical relationships between the samples. Finally, clustered results were visualized using clustermap function in seaborn [76].

#### Sequence logo analysis

To identify sequence features within each cluster, we employed a thresholding approach based on the saliency scores. Specifically, for each cluster, we computed the frequency of each amino acid for residues with a saliency score >0.5. Then, sequence logos were generated by Logomaker in Python [77].

#### Structural analysis of saliency score

For those HA stem antibodies with structural information available, the relationship between saliency score of each residue and its minimum distance to HA was examined. Distance was calculated using the application programming interface in PyMOL (Schrödinger).

### Mammalian cell culture

HEK293T cells were cultured in Dulbecco’s modified Eagle’s medium (DMEM high glucose; Gibco) supplemented with 10% heat-inactivated fetal bovine serum (FBS; Gibco), 1% penicillin-streptomycin (Gibco), and 1× GlutaMax (Gibco). Cell passaging was performed every 3 to 4 days using 0.05% Trypsin-EDTA solution (Gibco). Expi293F cells were maintained in Expi293 Expression Medium (Thermo Fisher Scientific). Sf9 cells (*Spodoptera frugiperda* ovarian cells, female, ATCC) were maintained in Sf-900 II SFM medium (Thermo Fisher Scientific).

### Expression and purification of mini-HA and HA

The mini-HA #4900 [58] and H1N1 A/Solomon Island/3/2006 HA were fused with N-terminal gp67 signal peptide and a C-terminal BirA biotinylation site, thrombin cleavage site, trimerization domain, and a 6xHis-tag, and then cloned into a customized baculovirus transfer vector [46]. Subsequently, recombinant bacmid DNA was generated using the Bac-to-Bac system (Thermo Fisher Scientific) according to the manufacturer’s instructions. Baculovirus was generated by transfecting the purified bacmid DNA into adherent Sf9 cells using Cellfectin reagent (Thermo Fisher Scientific) according to the manufacturer’s instructions. The baculovirus was further amplified by passaging in adherent Sf9 cells at a multiplicity of infection (MOI) of 1. Recombinant mini-HA protein was expressed by infecting 1 L of suspension Sf9 cells at an MOI of 1. On day 3 post-infection, Sf9 cells were pelleted by centrifugation at 4000 × *g* for 25 min, and soluble recombinant mini-HA and HA were purified from the supernatant by affinity chromatography using Ni Sepharose excel resin (Cytiva) and then size exclusion chromatography using a HiLoad 16/100 Superdex 200 prep grade column (Cytiva) in 20 mM Tris-HCl pH 8.0, 100 mM NaCl. The purified mini-HA protein was concentrated by Amicon spin filter (Millipore Sigma) and filtered by 0.22 µm centrifuge tube filters (Costar). Concentration of the protein was determined by nanodrop (Fisher Scientific). Proteins were subsequent aliquoted, flash frozen by dry-ice ethanol mixture, and stored at -80°C until used.

### Expression and purification of IgG

The heavy and light chain genes of the obtained antibody were synthesized as eBlocks (Integrated DNA Technologies), and then cloned into human IgG1 and human kappa or lambda light chain expression vectors using Gibson assembly according to a previously described method [78]. The plasmids were transiently co-transfected into HEK293T cells at a mass ratio of 2:1 (HC:LC) using Lipofectamine 2000 (Thermo Fisher Scientific). On day 3 post-transfection, supernatant containing the IgG was collected for binding experiment. The expression of IgG was confirmed by SDS-PAGE gel electrophoresis and Coomassie Blue R-250 staining. Selected IgGs were purified using a CaptureSelect CH1-XL Pre-packed Column (Thermo Fisher Scientific).

### Expression and purification of Fab

Fab heavy and light chains were cloned into phCMV3 vector. The plasmids were transiently co-transfected into Expi293F cells at a mass ratio of 2:1 (HC:LC) using ExpiFectamine 293 Reagent (Thermo Fisher Scientific). After transfection, the cell culture supernatant was collected at 6 days post-transfection. The Fab was then purified using a CaptureSelect CH1-XL pre-packed column (Thermo Fisher Scientific).

### Enzyme-linked immunosorbent assay (ELISA)

Nunc MaxiSorp ELISA plates (Thermo Fisher Scientific) were utilized and coated with 100 μL of recombinant proteins at a concentration of 1 μg ml^-1^ in a 1× PBS solution. The coating process was performed overnight at 4°C. On the following day, the ELISA plates were washed three times with 1× PBS supplemented with 0.05% Tween 20, and then blocked using 200 μL of 1× PBS with 5% non-fat milk powder for 2 hours at room temperature. After the blocking step, 100 μL of IgGs from the supernatant were added to each well and incubated for 2 hours at 37°C. The ELISA plates were washed three times to remove any unbound IgGs. Next, the ELISA plates were incubated with horseradish peroxidase (HRP)-conjugated goat anti-human IgG antibody (1:5000, Invitrogen) for 1 hour at 37°C. Subsequently, the ELISA plates were washed five times using PBS containing 0.05% Tween 20. Then, 100 μL of 1-Step Ultra TMB-ELISA Substrate Solution (Thermo Fisher Scientific) was added to each well. After 15 min incubation, 50 μL of 2 M H_2_SO_4_ solution was added to each well. The absorbance of each well was measured at a wavelength of 450 nm using a Sunrise absorbance microplate reader (BioTek Synergy HTX Multimode Reader).

### Biolayer interferometry binding assay

Binding assays were performed by biolayer interferometry (BLI) using an Octet Red96e instrument (FortéBio) at room temperature as described previously [79]. Briefly, His-tagged mini-HA proteins at 0.5 μM in 1× kinetics buffer (1× PBS, pH 7.4, 0.01% w/v BSA and 0.002% v/v Tween 20) were loaded onto anti-Penta-HIS (HIS1K) biosensors and incubated with the indicated concentrations of Fabs. The assay consisted of five steps: (1) baseline: 60 s with 1× kinetics buffer; (2) loading: 60 s with His-tagged mini-HA proteins; (3) baseline: 60 s with 1× kinetics buffer; (4) association: 60 s with Fab samples; and (5) dissociation: 60 s with 1× kinetics buffer. For estimating the exact *K*_D_, a 1:1 binding model was used.

### Virus neutralization assay

Madin-Darby canine kidney (MDCK) cells were seeded in a 96-well, flat-bottom cell culture plate (Thermo Fisher). The next day, serially diluted monoclonal antibodies were mixed with an equal volume of virus and incubated at 37°C for 1 hour. The antibody/virus mixture was then incubated with the MDCK cells at 37°C after the cells were washed twice with PBS. Following a 1-hour incubation, the antibody/virus mixture was replaced with Minimum Essential Medium (MEM) supplemented with 25 mM of 4-(2-hydroxyethyl)-1-piperazineethanesulfonic acid (HEPES) and 1 μg mL^-1^ of Tosyl phenylalanyl chloromethyl ketone (TPCK)-trypsin. The plate was incubated at 37°C for 72 hours and the presence of virus was detected by hemagglutination assay. The results were analyzed using Prism software (GraphPad).

### Cryogenic electron microscopy (cryo-EM) analysis

To prepare cryoEM grid, an aliquot of 4 μL purified protein at ∼0.5 mg mL^-1^ concentration with 7.5 μM lauryl maltose neopentyl glycol (LMNG) was applied to a 200-mesh Quantifoil 2Um Cu grid that was pre-treated with glow-discharge. Subsequently, the grid was blotted in a Vitrobot Mark IV machine (force = 0, time = 3 seconds), and plunge-frozen in liquid ethane. The grid was then loaded in a ThermoFisher Glacios microscope with a Volta Phase Plate and Falcon4 Direct Electron Detector. Data collection was done with Smart EPU software. Images were recorded at 130,000× magnification, corresponding to a pixel size of 0.96 Å/pix at super-resolution mode of the camera. A defocus range of -0.6 μm to -3 μm was set. A total dose of 52.76 e^−^/Å^2^ of each exposure was fractionated into 40 frames. CryoEM data processing was performed with cryoSPARC v4.3.0 following regular single-particle procedures. The CryoEM experiment was performed at the UIUC Materials Research Laboratory Central Research Facilities. Statistics are provided in **Table S4**. Structure was visualized using UCSF ChimeraX v1.5 (UCSF).

## Data availability

The cryoEM map of 310-18A5 Fab in complex with SI06 HA can be accessed at the Electron Microscopy Data Bank (EMDB) using accession code EMD-41849.

## Model and code availability

Custom python scripts for all analyses and model training have been deposited to: https://github.com/nicwulab/HA_Abs.

