## Supplementary material for "An explainable language model for antibody specificity prediction using curated influenza hemagglutinin antibodies": Figures S1-S7 and Table S4

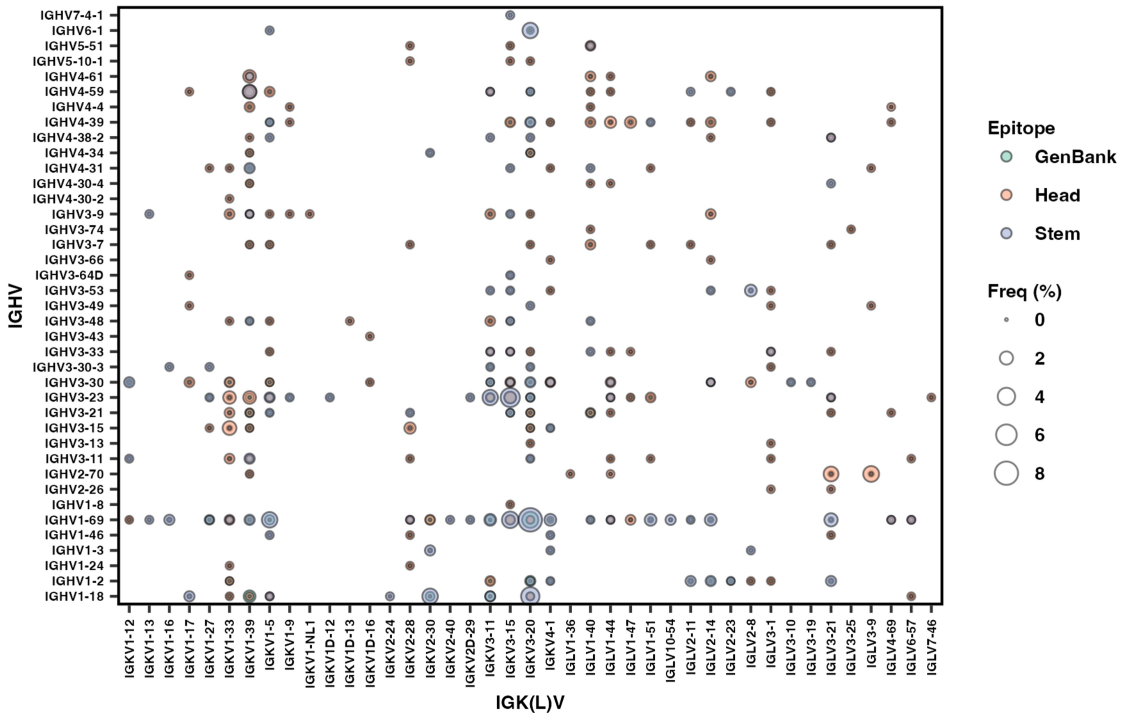


**Supplementary Figure 1. Preference of V gene pairings in influenza HA antibodies.** The frequencies of different V gene pairings between heavy and light chains are shown for influenza HA antibodies to the head and stem domains. Antibodies from GenBank were also included as a reference. The size of each data point represents the frequency of the corresponding IGHV/IGK(L)V pair within its specificity category. Only those antibodies with both IGHV and IGK(L)V gene information available were included in this analysis.

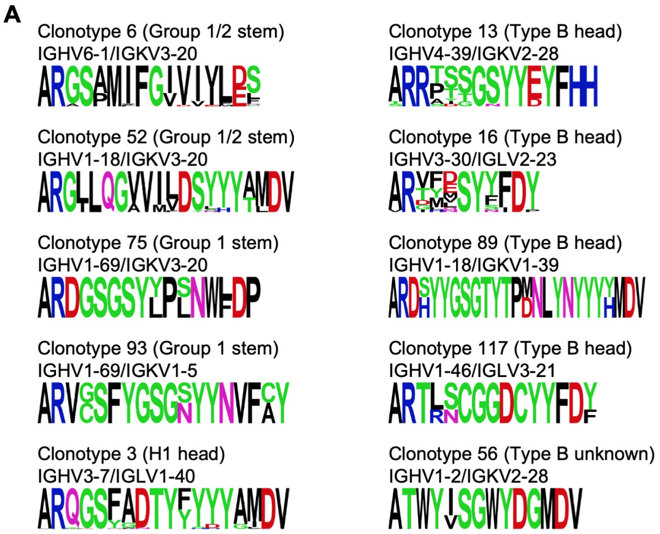


**B**

028-10134_4B04 (Henry et al. 2019)
Donor: male, age 37 (USA)

```
 C  A  R  D  H  Y  Y  G  S  G  T  Y  T  P  D  N  L  Y  N  Y  Y  Y  Y  M  D  V  W
TGTGCGCGCGATCATTACTATGGTTCGGGGACATACACACCGGACAATTTGTACAACTACTACTACTACATGGACGTCTGG
TGTGCGAGAGA  ATTACTATGGTTCGGGGA                    TACTACTACTACTACTACATGGACGTCTGG
IGHV1-18*04     IGHD3-10*01                                   IGHJ6*03

Total gene-derived nucleotides: 53
Total non-gene-derived nucleotides: 22
```

K77-2D11 (Liu et al. 2019)
Donor: male, age 38 (Australia)

```
 C  A  R  D  S  Y  Y  G  S  G  T  Y  T  P  M  N  L  Y  N  Y  Y  Y  H  M  D  V  W
TGTGCGAGAGATTCTTACTATGGTTCGGGGACTTATACTCCAATGAACCTTTACAACTATTATTACCATATGGACGTCTGG
TGTGCGAGAGA   TTACTATGGTTCGGGGAGTTAT              TTACTACTACTACTACGGTATGGACGTCTGG
IGHV1-18*01        IGHD3-10*01                                IGHJ6*02

Total gene-derived nucleotides: 58
Total non-gene-derived nucleotides: 17
```

**Supplementary Figure 2. Public clonotypes of influenza HA antibodies. (A)** Antibodies with the same IGHV/IGK(L)V genes and at least 80% sequence identity in the CDR H3 were defined as a clonotype. A clonotype with antibodies from at least two donors was defined as a public clonotype. A total of 10 public clonotypes were identified. The V gene usage and CDR H3 sequence are shown for each of these 10 public clonotypes. The CDR H3 sequences are shown as a sequence logo, where the height of each letter represents the frequency of the corresponding amino-acid variant (single-letter amino acid code) at the indicated position. This analysis captured many known recurring sequence features in HA stem antibodies, including IGHV6-1 with an [I/V]FG[I/L/V] motif (clonotype 6) [1, 2], VH1-18 with a QxxV motif in CDR H3 (clonotype 52) [3], and IGHV1-69 with a Tyr in the CDR H3 (clonotypes 75 and 93) [4]. The recurring usage of

IGHV3-7/IGLV1-40 among HA head antibodies (clonotype 3) was also noted previously [5, 6]. **(B)** Among the five public clonotypes to influenza type B HA, clonotypes 13, 16, 56, and 117 consisted of antibodies from the same study (**Table S1**) [7, 8]. In contrast, clonotype 89 consisted of antibodies from two different studies [8, 9]. Our dataset contained two antibodies within clonotype 89, namely 028-10134_4B04 and K77-2D11, which were isolated from donors in the US and Australia, respectively [8, 9]. Amino acid and nucleotide sequences of the V-D-J junction are shown for 028-10134_4B04 and K77-2D11. Antibody 028-10134_4B04 was isolated from a 37-year-old male in the US [8], whereas K77-2D11 was isolated from a 38-year-old male in Australia [7]. Putative germline sequences and segments were identified by IgBlast [10] and are indicated. Somatically mutated nucleotides are underlined. Intervening spaces at the V-D and D-J junctions are N-nucleotide additions. Both 028-10134_4B04 and K77-2D11 have a long CDR H3 with 25 amino acids (IMGT numbering), including a YYGSGTY that is largely encoded by IGHD3-10 and a TPxNL motif that is encoded by N-nucleotide addition. While previous studies of recurring sequence features among HA antibodies have mainly focused on influenza type A HA [1-4, 11-15], our results suggest that recurring sequence features among antibodies to influenza type B HA may also be quite common.

**A**

```
IGHD4-17: TGACTACGGTGACTAC
Frame 1:   * L R * L
Frame 2:    D Y G D Y
Frame 3:     T T V T
```

**B**

```
W85-1A07 (IGHV3-48/IGLV2-14)
 C  A  R  A  H  M  I  Y  G  D  H  V  H  L  N  A  F  D  I  W
TGTGCGAGAGCTCATATGATATACGGTGACCACGTTCATCTGAATGCTTTTGATATTTGG

019_10_4A03 (IGHV3-15/IGKV2-28)
 C  T  T  D  Y  G  D  Y  L  N  G  G  R  W
TGTACCACTGACTACGGTGACTACCTTAACGGGGGGCGCTGG

150055-029_3E03 (IGHV3-23/IGKV1-33)
 C  A  K  G  G  Y  G  D  N  G  L  D  V  F  D  I  W
TGTGCGAAAGGAGGATACGGTGACAACGGGTTGGATGTCTTTGATATCTGG

SFV009_3F05 (IGHV3-7/IGKV1-5)
 C  A  R  A  G  S  Y  G  D  Y  R  P  I  N  N  W  F  D  P  W
TGTGCGAGAGCGGGGAGTTACGGTGACTACAGGCCGATAAACAACTGGTTCGACCCCTGG

SFV019_2A02 (IGHV1-18/IGKV1-33)
 C  A  R  R  G  D  Y  G  D  Y  R  G  D  A  F  D  I  W
TGTGCGAGACGTGGGGACTACGGTGACTACCGGGGTGATGCATTTGATATCTGG
```

**Supplementary Figure 3. YGD motif in IGHD4-17 HA head antibodies. (A)** The nucleotide sequence of IGHD4-17 and its amino acid sequences in all three translation frames are shown. **(B)** CDR H3 sequences of representative IGHD4-17 HA head antibodies with different V gene usages. The YGD motif and the IGHD4-17-encoded region are highlighted in red.

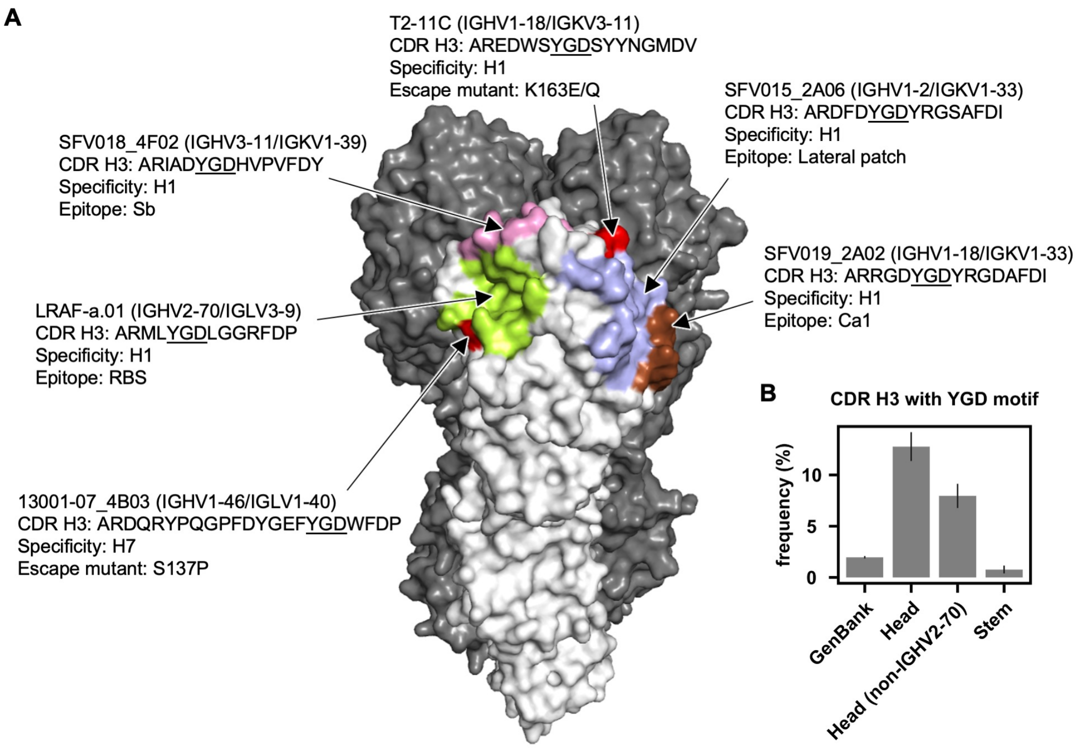


**Supplementary Figure 4. Representative IGHD4-17 HA head antibodies with YGD motif. (A)** Different epitopes, including the receptor-binding site (RBS, lime), Sb (pink), Ca1 (brown), and lateral patch (blue), are shown on the HA structure (PDB: 3LZG) [16]. Information of representative IGHD4-17 HA head antibodies that target these epitopes is shown. The locations of K163E/Q and S137P, which escape antibodies T2-11C [6] and 13001-07_4B03 [17], respectively, are colored in red. Of note, S137P (H3 numbering) was named as S152P in the original paper [17]. **(B)** Frequency of antibodies with a YGD motif in the CDR H3 among all antibodies from Genbank, HA head antibodies, non-IGHV2-70-encoded HA head antibodies, and HA stem antibodies.

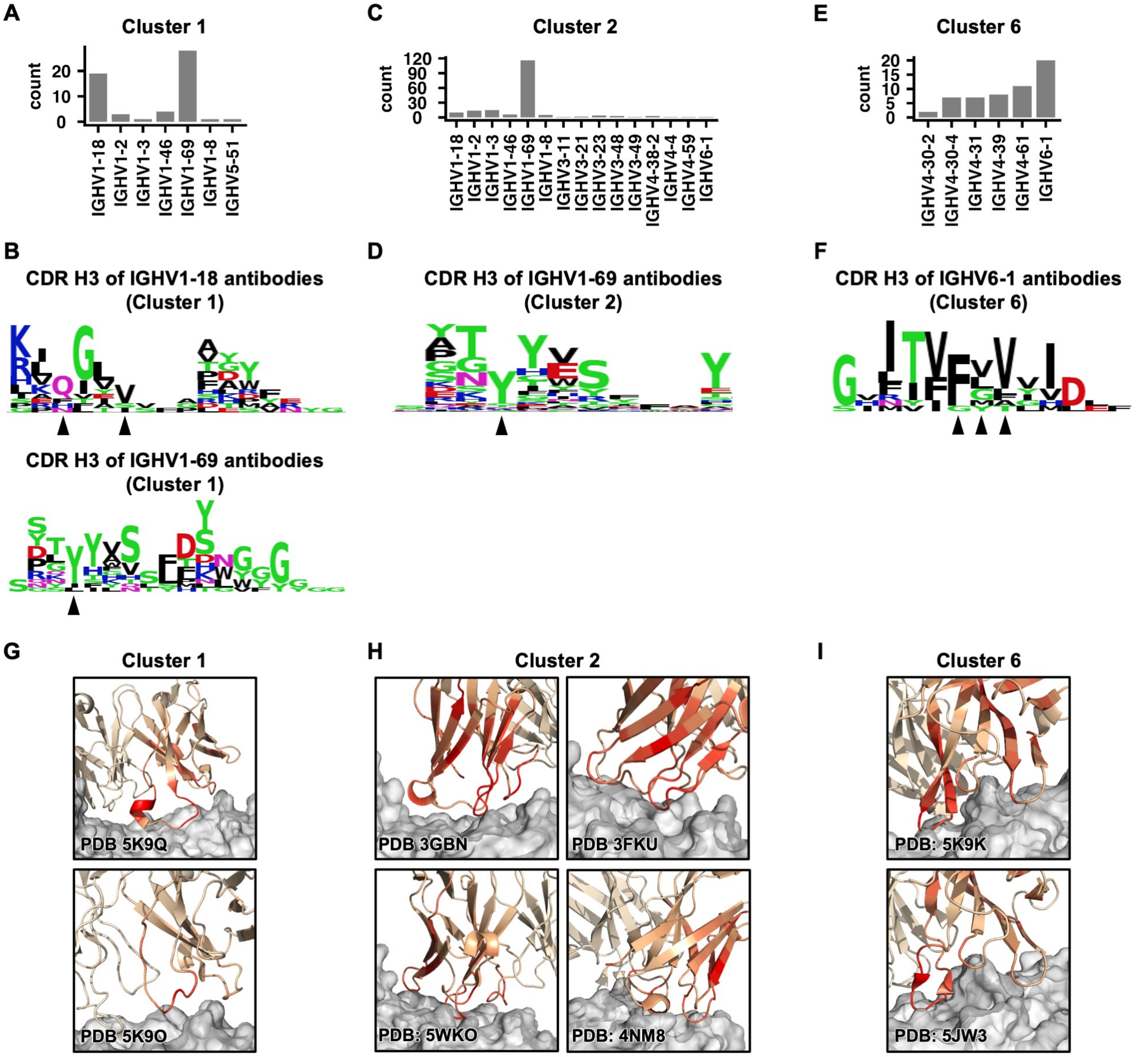


**Supplementary Figure 5. Sequence features of clusters 1, 2, and 6 of HA stem antibodies.** **(A, C, E)** IGHV gene usages among antibodies in **(A)** cluster 1, **(C)** cluster 2, and **(E)** cluster 6 are shown **(B, D, F)** The saliency score of each CDR H3 residue in **(B)** IGHV1-18 antibodies (top) and IGHV1-69 antibodies (bottom) within cluster 1, **(D)** IGHV1-69 antibodies within cluster 2, and **(F)** IGHV6-1 antibodies within cluster 6 was analyzed. The frequency of each amino acid for residues with a saliency score >0.5 is shown as a sequence logo. Arrows at the bottom indicate the residues of interest, including **(B)** a QxxV motif (top), Y98 (bottom), **(D)** Y98, and **(F)** an FGV motif **(G-I)** Saliency scores are projected onto the structures of **(G)** two antibodies in cluster 1

(PDB 5K9Q and PDB 5K9O [3]), **(H)** four antibodies in cluster 2 (PDB 3GBN [18], PDB 3FKU [19], PDB 5WKO [11], and PDB 4NM8 [20]), and **(I)** two antibodies in cluster 6 (PDB 5K9K [3] and PDB 5JW3 [21]). Color scheme is the same as that in **Figure 4A**.

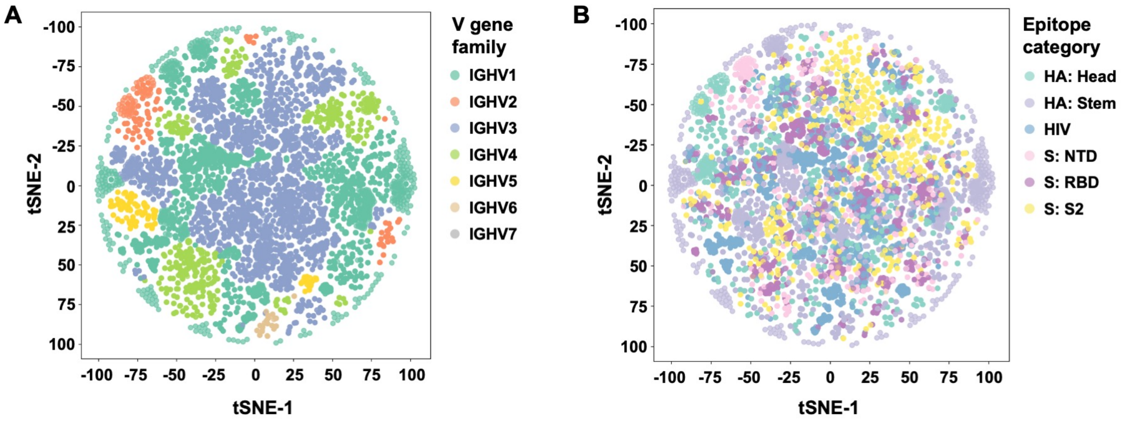


**Supplementary Figure 6. t-SNE analysis of the final-layer embeddings of the pre-trained mBLM.** The final-layer embeddings of the pre-trained mBLM model (i.e. prior to fine-tuning for specificity prediction) was analyzed by t-SNE (t-distributed Stochastic Neighbor Embedding). Heavy chain sequences in the training set for fine-tuning were used in this analysis. Each datapoint represents one heavy chain sequence. Datapoints are colored by **(A)** V gene families or **(B)** specificity categories.

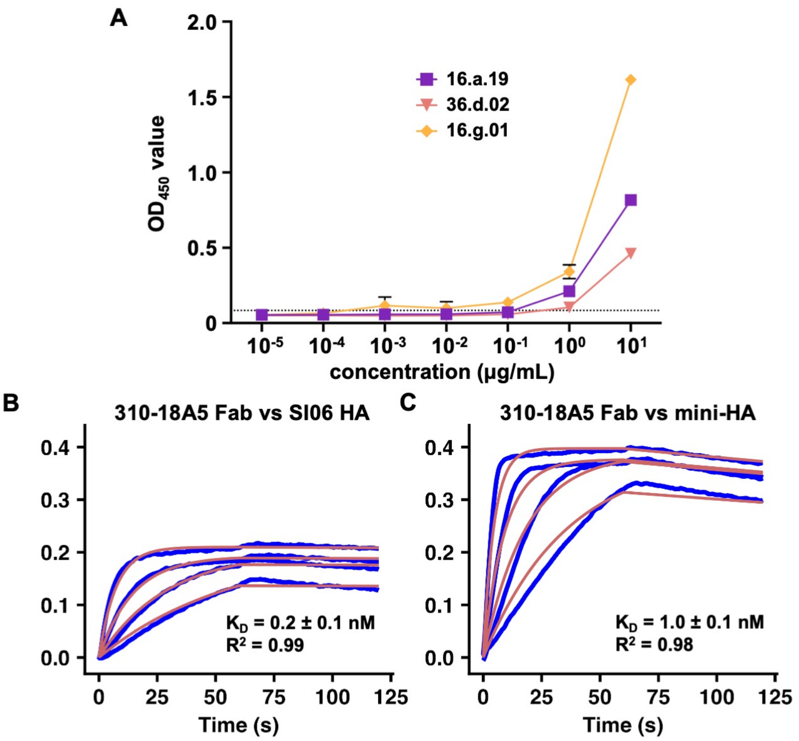


D

| Strain | MIC (μg/mL) |
|---|---|
| H1N1 A/Puerto Rico/8/1934 | 18.75 |
| H1N1 A/Michigan/45/2015 | 37.5 |

**Supplementary Figure 7. Binding and neutralization activity of antibodies that were predicted to target HA stem. (A)** ELISA was used to test the binding of purified antibodies 16.a.19, 36.d.02, and 16.g.01 at different concentrations to mini-HA. PBS was used as a negative control (dotted line). Data are representative of two independent experiments. **(B-C)** Binding kinetics of 310-18A5 Fab against **(B)** H1N1 A/Solomon Islands/3/2006 (SI06) HA and **(C)** mini-HA were measured by biolayer interferometry (BLI). Y-axis represents the response. Blue lines represent the response curve and red lines represent the 1:1 binding model. Binding kinetics were measured for four concentrations of Fab at 2-fold dilution ranging from 200 nM to 25 nM. Dissociation constant ($K_D$) and the goodness of model fitting ($R^2$) are indicated. **(D)** Neutralization activity of 310-18A5 was tested against two H1N1 strains, namely A/Puerto Rico/8/1934 and A/Michigan/45/2015. Minimal inhibitory concentration (MIC) is indicated.

**Table S4. Cryo-EM data collection statistics.**

| | SI06HA-18A5 complex (EMD-41849) |
|---|---|
| **Data collection and processing** | |
| Magnification | 130,000 |
| Voltage (kV) | 200 |
| Electron exposure (e-/Å2) | 52.76 |
| Defocus range (μm) | -0.6 to -3 |
| Pixel size (Å) | 0.96 |
| Symmetry imposed | C3 |
| Initial particle images (no.) | 41,774 |
| Final particle images (no.) | 39,446 |
| Map resolution (Å) | 4.81 |
| FSC threshold | 0.143 |
| Map resolution range (Å) | N/A |
